# Structural analysis of human papillomavirus E6 interactions with Scribble PDZ domains

**DOI:** 10.1101/2022.10.12.511839

**Authors:** Bryce Z. Stewart, Sofia Caria, Patrick O. Humbert, Marc Kvansakul

## Abstract

The cell polarity regulator Scribble has been shown to be a critical regulator of the establishment and development of tissue architecture, and its dysregulation promotes or suppresses tumor development in a context dependent manner. Scribble activity is subverted by numerous viruses. This includes human papillomaviruses (HPVs), who target Scribble via the E6 protein. Binding of E6 from high-risk HPV strains to Scribble via a C-terminal PDZ binding motif leads to Scribble degradation *in vivo*. However, the precise molecular basis for Scribble-E6 interactions remains to be defined. We now show that Scribble PDZ1 and PDZ3 are the major interactors of HPV E6 from multiple high-risk strains, with each E6 protein displaying a unique interaction profile. We then determined crystal structures of Scribble PDZ1 and PDZ3 domains in complex with the PBM motifs of E6 from HPV strains 16, 18 and 66. Our findings reveal distinct interaction patterns for each E6 PBM motif from a given HPV strain, suggesting that a complex molecular interplay exists that underpins the overt Scribble-HPV E6 interaction and controls E6 carcinogenic potential.

## INTRODUCTION

Cell polarity is a critical feature of eukaryotic cells and plays a crucial role in tissue development and correct establishment of tissue architecture [1]. At the molecular level, cell polarity leads to the asymmetric distribution of macromolecules including proteins, lipids and carbohydrates into distinct cellular domains, which gives rise to apical-basal cell polarity [2]. Apical-basal cell polarity plays a significant role in the regulation of important cellular signalling pathways including those associated with apoptosis, vesicle trafficking, cell proliferation and migration [3]. Importantly, loss of cell polarity is an important hallmark of cancer development and viral infection [4]. Apical-basal cell polarity is controlled by the dynamic interplay between three polarity modules, Scribble, Par and Crumbs [5]. Scribble is a key regulator of cell polarity and a component of the Scribble module comprising Scribble, Discs-large (Dlg) and Lethal-Giant-Larvae (Lgl). After its initial discovery in *Drosophila melanogaster* as a tumour suppressor [6], Scribble has been shown to be play a role in tumour initiation, and when paired with oncogenic drivers such as RAS, its loss can drive tumour progression in multiple epithelial tissue types including mammary, prostate, skin and the lung [7]. In addition to its tumour modulatory activities, Scribble regulates polarity and signalling in a wide number of cell types and organisms, playing a crucial role in organ development and physiology [8].

Scribble features 16 Leucine Rich Repeats and 4 PSD-95/Disc-large/ZO-1 (PDZ) domains, and is a member of the LAP family of proteins [7]. Being a signalling adaptor protein, the vast majority of Scribble interactions are mediated via its PDZ domains, allowing Scribble to function within multiple discrete signalling pathways [3]. Interactions with Scribble PDZ domains are mediated by PDZ binding motifs (PBMs), short amino acid sequences often found at the very C-termini of interacting proteins. PBMs are divided into three classes, with class 1 PBMs consisting of X-T/S-X-ϕ_COOH_ (where X is any residue and ϕ is any hydrophobic residue), class 2 being X-ϕ-X-ϕ _COOH_ and class 3 is X-D/E-X-ϕ _COOH_ [9–11].

Cell polarity signalling is subverted by a range of viruses [4] in order to support viral infection and proliferation, including SARS-CoV-2 [12], HTLV [13], TBEV [14], influenza virus [15] and HPVs, where viral effector proteins engage with components of cell polarity components including Scribble, Pals1 or Dlg.

HPVs commonly cause plantar warts that, after a time, rescind. However, several strains of HPV are found to be carcinogenic [16]. Almost all cervical cancers are associated with HPV and women carrying the virus are up to 50 times more likely to develop cervical cancer than those without [17, 18]. HPV strains 16 and 18 are found to cause almost 90% of associated cancers, with HPV-16 being the most prevalent [19]. While the virus is common amongst women, few develop cancer. It is now well established that oncogenic strains that persist are key to cervical cancer development [20], and a range of HPV strains including 16 and 18, along with 31 and 33, have been declared oncogenic by the world health organisation (WHO) [21]. HPV encodes for E6, a viral oncoproteins, that interacts with key host proteins that regulate apoptosis and polarity, including the cell polarity regulating tumour suppressing proteins human Scribble and Dlg1 [22, 23]. This interaction is mediated via a PDZ binding motif (PBM) at the extreme C-termini of E6 from high risk E6 oncoproteins [24, 25]. E6 subsequently associates with host ubiquitin ligases and any protein that interact with HPV E6 are subsequently degraded [26].

HPV E6 has been shown to bind to many protein interactors including p53, Scribble and Dlg [23, 27]. It was shown that Scribble is the only protein that directly regulates E6 expression levels by transcription and translation, and moreover, Scribble was also identified as the tightest binder amongst these [22, 27, 28]. Scribble has been shown to bind to E6 originating from oncogenic HPV strains only, which feature mostly class 1 PBMs, but no interaction has been observed with E6 from non-oncogenic strains such as HPV 40 [27]. This is contrast to human Dlg1 which is a common target of the HPV E6 PBM regardless of transforming potential [27]. E6 from HPV16 has been shown to bind indiscriminately to the tandem PDZ1/2 and PDZ3/4 domains of human Scribble [29]. However, subsequent pull-down experiments suggested that E6 from HPV16 and 18 primarily engage the Scribble PDZ3 domain [25]. Whether or not E6 from different HPV strains harbor particular patterns of interaction with specific PDZ domains of Scribble, and the exact nature as to why Scribble PDZ domains may display selectivity for E6 is poorly understood. We have performed a detailed analysis of Scribble PDZ domain interactions with 8-mer peptides of the C-termini HPV E6 oncogenic strains 16, 18, 31, 33, 51, 66 and 70 as well as non-oncogenic strain 40. We then determined crystal structures of Scribble PDZ domains bound to interacting E6 PBM motifs. Our studies show that Scribble PDZ domains 1 and 3 both bind tightly, as compared to other Scribble PDZ domain interactions, to all oncogenic strains and the highest average affinity for E6 16. Intriguingly, our data also shows weak binding between Scribble PDZ 2 domain and most oncogenic HPV E6 strains except E6 16 and 66.

## RESULTS

### Isolated Scribble PDZ domains specifically interact with the high risk HPV E6 PBM

Scribble has previously been shown to directly interact with high-risk HPV16 and HPV18 E6 C-terminal PDZ binding motif (PBM) via its PDZ domains using pull-down assays. Furthermore, E6 from all cancer-causing HPV strains bound to Scribble, including E6 from HPV 31 and 33 [27]. These data suggested that Scribble binding affinity may be a good oncogenic predictor. To further understand these interactions, we examined the affinity of recombinant isolated Scribble PDZ1, 2, 3 and 4 domains for a panel of 10-mer peptides of E6 corresponding to the major oncogenic HPV strains (Table 1). Our ITC measurements revealed a distinct pattern of interaction for different HPV E6 peptides that interact with Scribble. As expected, E6 from HPV 40 did not bind to any PDZ domain. Whilst all E6 peptides that interacted with Scribble bound PDZ1 and 3 domains, we also observed binding of E6 from HPV 18, 31, 33, 51 and 70 to PDZ2. In contrast, E6 from HPV 16 and 66 did not bind to PDZ2. E6 from HPV 16 bound PDZ1 and 3 with similar affinity, whereas E6 from HPV 31, 51, 66 and 70 bound PDZ1 more tightly than PDZ3. E6 from HPV 18 and 33 bound PDZ3 more tightly than PDZ1. PDZ2 interactions with E6 were in all cases the weakest interaction measured for a given HPV strain. Interestingly, our measurements reveal binding of HPV60 E6 to Scribble protein for the first time, in contrast to previous reports.

**Table 1:** Summary of affinity and thermodynamic binding parameters for Scrib PDZ domains interactions with HPV E6 peptides. NB denotes no binding, n.d. denotes not determined. Each of the value was calculated from at least three independent experiments.

|  | Scrib PDZ1 |  |  |  | Scrib PDZ2 |  |  |  |
| --- | --- | --- | --- | --- | --- | --- | --- | --- |
| | $K_D$ (μM) | N | $-T\Delta S$ (cal/mol/K) | $\Delta H$ (kcal/mol) | $K_D$ (μM) | N | $-T\Delta S$ (cal/mol/K) | $\Delta H$ (kcal/mol) |
| HPV16 E6 PBM | $4.99 \pm 0.55$ | $1.11 \pm 0.01$ | $-2.05 \pm 0.36$ | $-3.62 \pm 0.43$ | NB | NB | NB | NB |
| HPV18 E6 PBM | $14.79 \pm 2.85$ | $1.07 \pm 0.03$ | $-3.95 \pm 0.43$ | $-1.64 \pm 0.54$ | $17.27 \pm 1.71$ | $1.13 \pm 0.06$ | $-4.90 \pm 0.14$ | $-1.07 \pm 0.11$ |
| HPV33 E6 PBM | $5.50 \pm 2.44$ | $1.10 \pm 0.08$ | $0.34 \pm 0.25$ | $-5.02 \pm 0.50$ | $21.84 \pm 1.86$ | $0.99 \pm 0.12$ | $-4.92 \pm 0.34$ | $-1.06 \pm 0.28$ |
| HPV31 E6 PBM | $6.97 \pm 1.64$ | $1.00 \pm 0.04$ | $-5.46 \pm 0.63$ | $-1.58 \pm 0.49$ | $12.32 \pm 0.93$ | $1.00 \pm 0.03$ | $-5.98 \pm 0.17$ | $-0.52 \pm 0.22$ |
| HPV51 E6 PBM | $7.73 \pm 1.02$ | $1.09 \pm 0.05$ | $-4.73 \pm 1.02$ | $-1.12 \pm 0.96$ | $17.30 \pm 2.90$ | $1.09 \pm 0.02$ | $-5.84 \pm 0.00$ | $-0.42 \pm 0.09$ |
| HPV66 E6 PBM | $9.27 \pm 0.69$ | $1.16 \pm 0.01$ | $-5.95 \pm 0.35$ | $-0.56 \pm 0.30$ | NB | NB | NB | NB |
| HPV70 E6 PBM | $5.14 \pm 0.31$ | $1.11 \pm 0.04$ | $-2.59 \pm 0.45$ | $-2.96 \pm 0.45$ | $14.11 \pm 2.64$ | $1.05 \pm 0.07$ | $-5.63 \pm 0.38$ | $-0.73 \pm 0.26$ |
| HPV40 E6 PBM | NB | NB | NB | NB | NB | NB | NB | NB |

|  | Scrib PDZ3 |  |  |  | Scrib PDZ4 |  |  |  |
| --- | --- | --- | --- | --- | --- | --- | --- | --- |
| | $K_D$ (μM) | N | $-T\Delta S$ (cal/mol/K) | $\Delta H$ (kcal/mol) | $K_D$ (μM) | N | $-T\Delta S$ (cal/mol/K) | $\Delta H$ (kcal/mol) |
| HPV16 E6 PBM | $4.41 \pm 1.66$ | $1.07 \pm 0.08$ | $4.22 \pm 0.78$ | $-11.55 \pm 0.54$ | NB | NB | NB | NB |
| HPV18 E6 PBM | $7.02 \pm 0.03$ | $1.13 \pm 0.08$ | $-1.41 \pm 0.99$ | $-5.62 \pm 0.98$ | NB | NB | NB | NB |
| HPV33 E6 PBM | $9.88 \pm 0.86$ | $1.09 \pm 0.09$ | $-0.55 \pm 0.14$ | $-6.28 \pm 0.09$ | NB | NB | NB | NB |
| HPV31 E6 PBM | $2.05 \pm 0.25$ | $1.15 \pm 0.03$ | $5.83 \pm 0.63$ | $-13.60 \pm 0.69$ | NB | NB | NB | NB |
| HPV51 E6 PBM | NB | NB | NB | NB | NB | NB | NB | NB |
| HPV66 E6 PBM | $11.51 \pm 1.39$ | $1.14 \pm 0.08$ | $-2.65 \pm 0.17$ | $-2.72 \pm 0.10$ | NB | NB | NB | NB |
| HPV70 E6 PBM | $14.23 \pm 3.21$ | $0.97 \pm 0.04$ | $-3.17 \pm 0.76$ | $-3.19 \pm 0.92$ | NB | NB | NB | NB |
| HPV40 E6 PBM | $10.96 \pm 0.88$ | $1.05 \pm 0.01$ | $1.03 \pm 0.83$ | $-7.79 \pm 0.88$ | NB | NB | NB | NB |

Examination of the thermodynamic binding parameters of HPV E6 peptides to Scribble PDZ1 (Figure 1, Table 1) revealed that E6 from HPV33 displays a different thermodynamic pattern compared to other PDZ1 interactors, by featuring an unfavourable entropy contribution. Interactions of E6 from HPV16, 33 and 70 with PDZ1 are driven primarily by enthalpic contributions, whereas E6 from HPV 18, 31, 51 and 66 are primarily entropy driven. In contrast, all PDZ3 interactions are more enthalpy driven. Notably, E6 from HPV 16, 33 and 70 all feature an unfavourable entropy contribution, which in case of HPV 16 and 33 is countered by a large favourable enthalpy. Interestingly, all PDZ2 interactions are dominated by entropic contributions.

**Figure 1.**
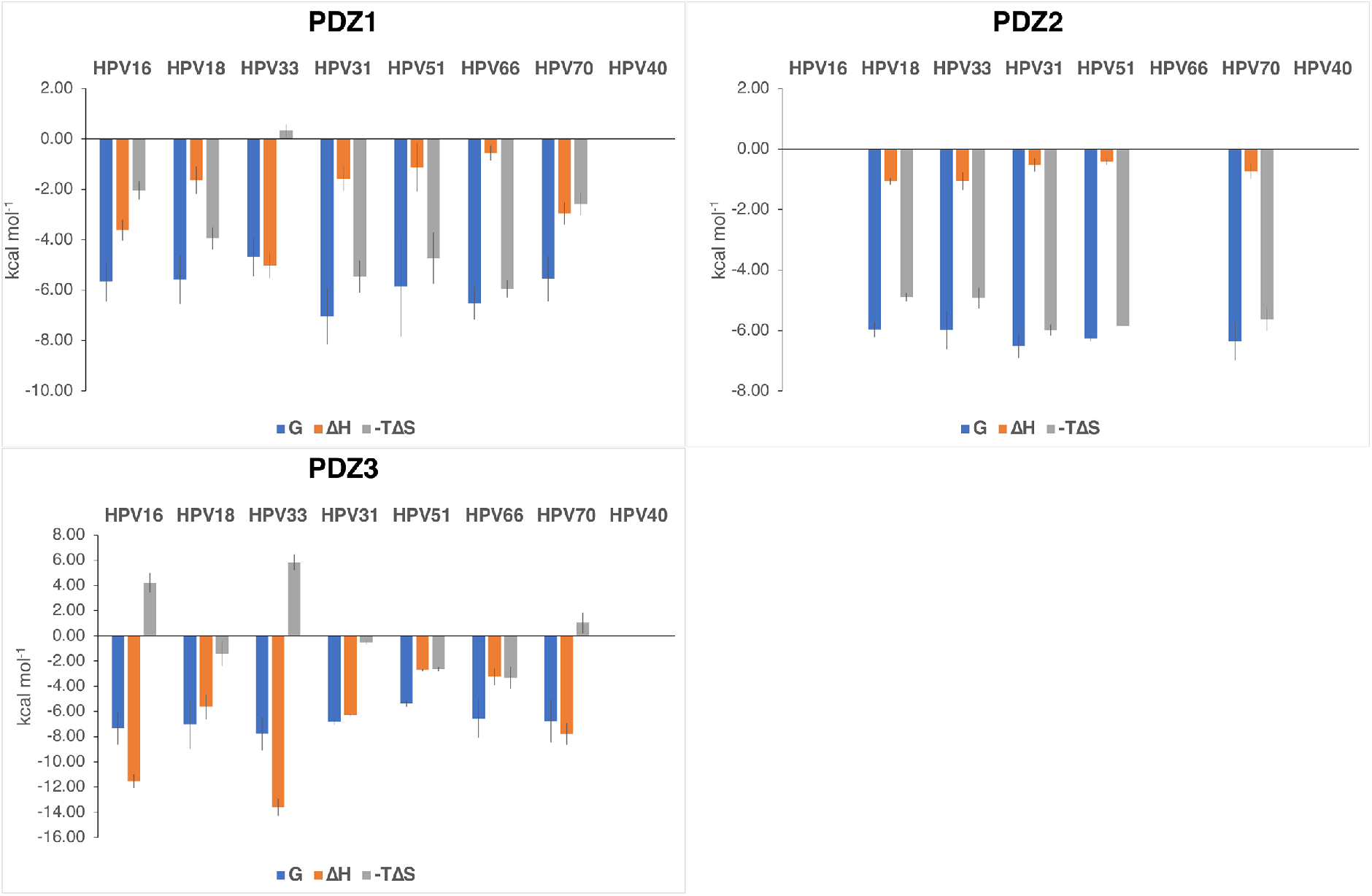
Thermodynamic analysis of Scribble PDZ domains 1, 2 and 3 binding to HPV E6 peptides. The enthalpy (-ΔH (kcal mol^−1^)) and entropy (-TΔS (kcal mol^−1^ K^−1^) of interactions between Scribble PDZ 1, 2 and 3 and HPV E6 PBM peptides are shown. Each value was calculated from at least three independent experiments. Peptides used are the C-terminal PBM motifs of HPV E6 strains 16, 18, 31, 33, 40, 51, 66 and 70.

### The crystal structures of PDZ:HPV E6 peptides

To understand the structural basis for the selectivity of each HPV E6 peptide we next determined the crystal structures of Scribble PDZ domains bound to certain HPV E6 PBM sequences (Figure 2, 3 and Table 2). Specifically, we determined structures of Scribble PDZ1 bound to E6 from HPV 16 and 18, as well as Scribble PDZ3 bound to HPV 16 and 66.

**Figure 2.**
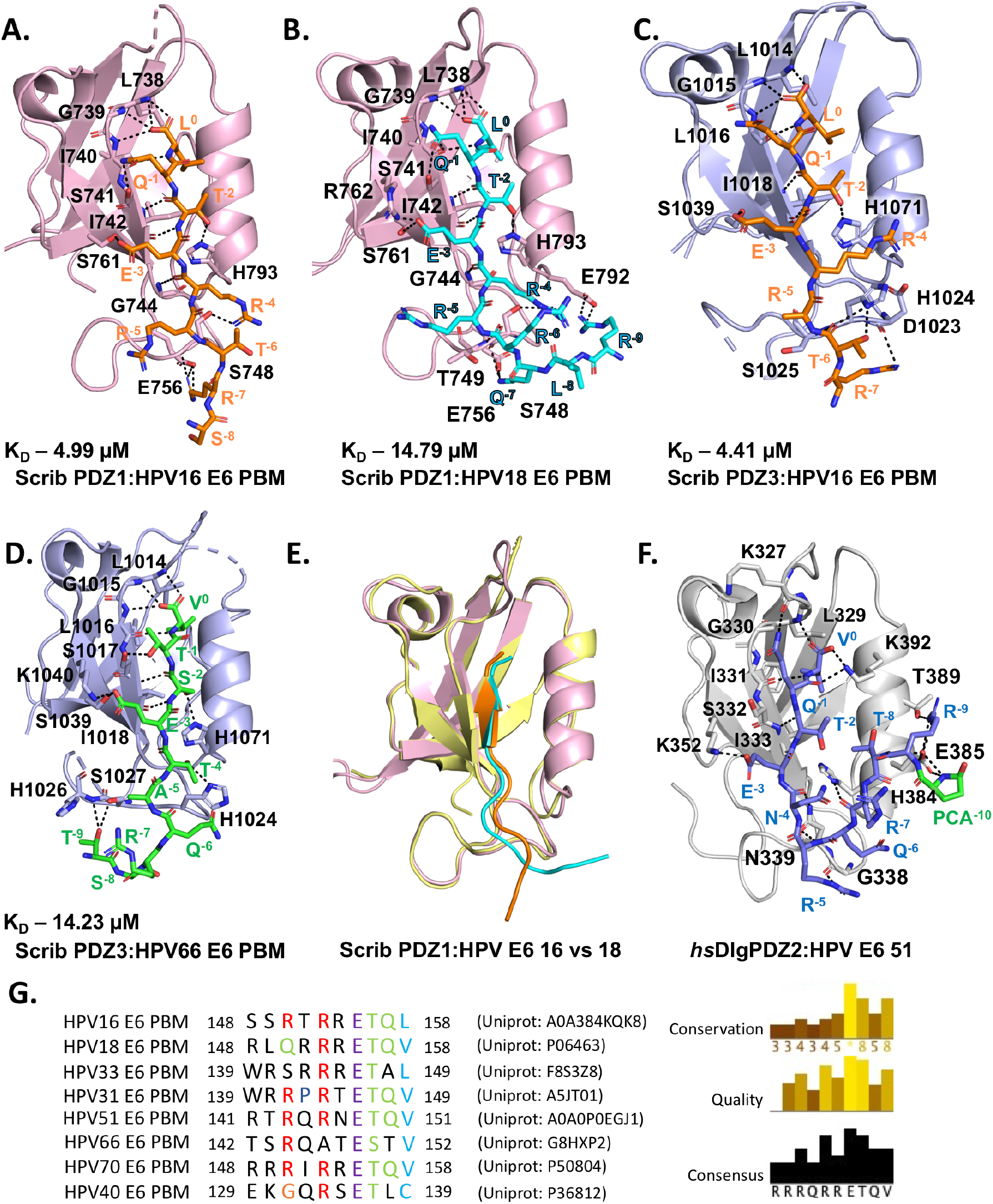
Interactions of Scribble PDZ domains with HPV E6 PBM. (**A**.) The crystal structures of PDZ1 bound to HPV E6 16, (**B**.), PDZ1 bound to HPV E6 18 (**C**.) PDZ3 bound to HPV E6 16 and (**D**.) PDZ3 bound to HPV E6 66. **E**. shows the superimposed structures of PDZ1:HPV E6 16 and 18. HPV E6 16, 18 and 66 peptides interact with the respective Scribble PDZ1 and PDZ3 domains via the canonical groove between β strand 2 and α helix 2. PDZ1 (pale yellow and light pink) is shown as cartoon with the respective HPV E6 16 peptide (cyan) and HPV E6 18 peptide (magenta) represented as sticks. PDZ3 (light blue) is shown as a cartoon with HPV E6 16 (orange) and HPV E6 66 (green) represented as sticks. **F**. shows *hs*Dlg1PDZ2 as cartoon (pale cyan) bound to HPV E6 51 in sticks (deep blue) with PDB accession ID: 2M3M. **G**. shows HPV E6 peptide sequence alignment. Uniprot accession numbers and peptide residue range are listed with corresponding peptides. Blue residues hydrophobic, magenta residues are negatively charged, red are positively charged, green are polar, yellow are prolines and white for unconserved. Three different conservation metrics calculated by Jalview are shown. Quality score is calculated for each column in an alignment by summing, for all mutations, the ratio of the two BLOSUM 62 scores for a mutation pair and each residue’s conserved BLOSUM62 score, and then plotted on a scale from 0 to 1. Displayed to the right of the alignment is the consensus percentage of the modal residue per column. The alignment was generated using Clustal Omega [51] and analysed using Jalview [52].

**Figure 3.**
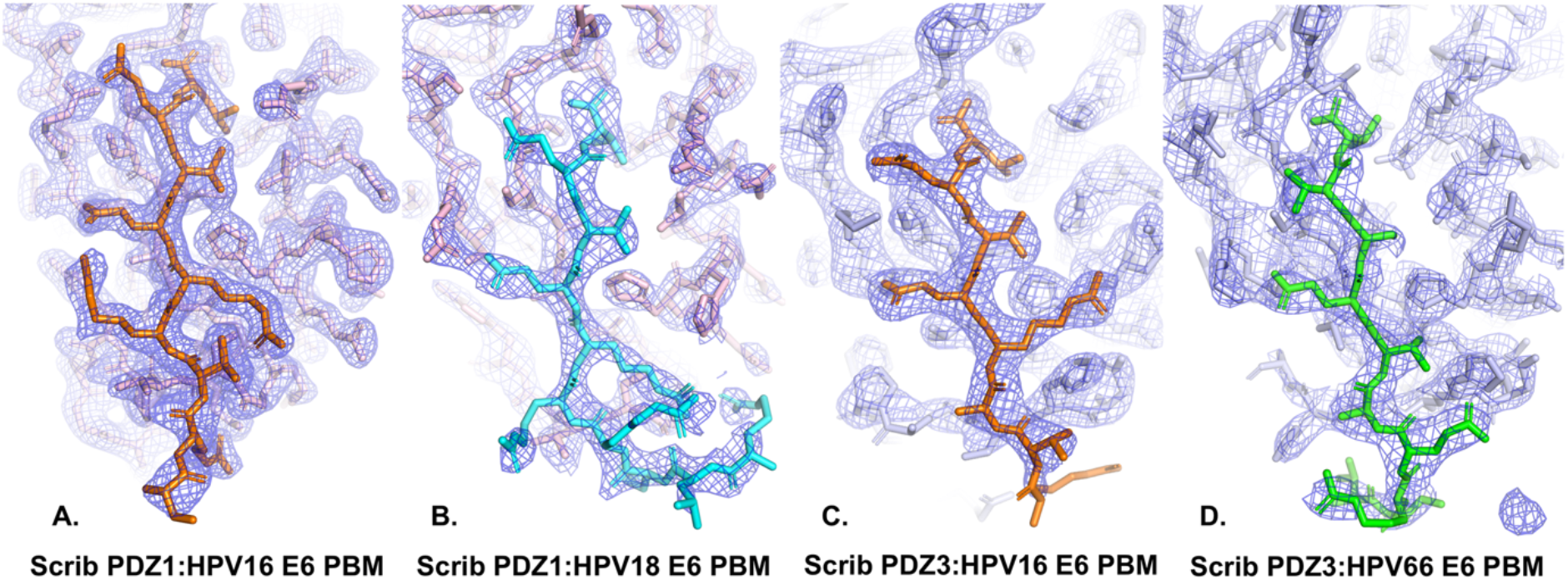
2Fo-Fc electron density maps of PDZ1 and PDZ3 complexes with peptides of HPV E6 strains 16, 18 and 66. Blue electron density map is shown as blue mesh and contoured at 1.5 σ, encompassing the binding groove of Scribble PDZ domains 1 (light pink) and 3 (pale blue) in complex with HPV E6 PBM peptides 16 (orange), 18 (cyan) and 66 (green) represented as sticks **A**. PDZ1:HPV E6 16, **B**. PDZ1:HPV E6 18, **C**. PDZ3:HPV E6 16 and **D**. PDZ3:HPV E6 66.

**Table 2:** X-ray data collection and refinement statistics.

|  | PDZ1:HPV E6 16 | PDZ1:HPV E6 18 | PDZ3:HPV E6 16 | PDZ3:HPV E6 66 |
| --- | --- | --- | --- | --- |
| **Data collection** |  |  |  |  |
| Space group | C 2 | P 2<sub>1</sub>2<sub>1</sub>2<sub>1</sub> | P 6<sub>1</sub>22 | C 2 |
| No of molecules in AU | 2+2 | 2+2 | 1+1 | 2+2 |
| Cell dimensions |  |  |  |  |
| *a*, *b*, *c* (Å) | 74.62, 31.14, 91.76 | 34.41, 55.25, 114.28 | 70.61, 70.61, 140.78 | 64.08, 90.00, 36.36 |
| $\alpha$, $\beta$, $\gamma$ (°) | 90.00, 107.56, 90.00 | 90.00, 90.00, 90.00 | 90.00, 90.00, 120.00 | 90.00, 91.92, 90.00 |
| Wavelength (Å) | 0.9537 | 0.9537 | 0.9537 | 0.9537 |
| Resolution (Å)* | 35.57-2.0 (2.07-2.0) | 39.72-2.8 (2.9-2.8) | 37.23-2.5 (2.59-2.5) | 32.02-2.90 (3.0-2.9) |
| *R*<sub>sym</sub> or *R*<sub>merge</sub>* | 0.079 (0.836) | 0.1381 (0.5239) | 0.1008 (1.363) | 0.0843 (0.8205) |
| *I* / $\sigma$ *I* * | 8.11 (1.17) | 12.18 (2.56) | 23.29 (1.61) | 6.16 (0.75) |
| CC(1/2) | 0.998 (0.54) | 0.997 (0.964) |  | 0.993 (0.44) |
| Completeness (%)* | 98.86 (99.5) | 98.23 (99.47) |  | 99.57 (99.56) |
| Redundancy* | 2.9 (2.9) | 6.2 (6.5) | 4.3 (6.2) | 1.9 (1.9) |
| Wilson B-factor | 33.63 | 47.23 | 65.73 | 72.87 |
| **Refinement** |  |  |  |  |
| Resolution (Å) | 35.57-2.0 | 39.72-2.8 | 37.23-2.5 | 32.02-2.9 |
| No. reflections | 13792 (1385) | 5770 (560) | 7707 (732) | 4616 (451) |
| *R*<sub>work</sub> / *R*<sub>free</sub> | 0.2194/0.2556 | 0.226/0.254 | 0.2741/0.2803 | 0.2696/0.2890 |
| No. non-hydrogen atoms |  |  |  |  |
| Protein | 1557 | 1777 | 737 | 1403 |
| Ligand/ion | 182 | 4 | 141 | 20 |
| Water | 82 | 66 | 26 | 11 |
| *B*-factors |  |  |  |  |
| Protein | 48.41 | 46.45 | 68.1 | 92.7 |
| Ligand/ion | 67.16 | 47.38 | 119.46 | 99.23 |
| Water | 47.38 | 42.62 | 74.16 | 87.37 |
| R.m.s. deviations |  |  |  |  |
| Bond lengths (Å) | 0.005 | 0.002 | 0.015 | 0.003 |
| Bond angle (°) | 0.52 | 0.47 | 1.62 | 0.67 |
| Ramachandran plot (%) |  |  |  |  |
| Favored | 98.96 | 98.67 | 95.56 | 93.68 |
| Allowed | 1.04 | 1.33 | 4.44 | 6.32 |
| Disallowed | 0.0 | 0.0 | 0.0 | 0.0 |

\* Values in parenthesis are for the highest resolution shell.

PDZ1 adopts a globular PDZ fold β-sandwich structure comprised of six β-strands and two α-helices as previously described [30]. HPV16 E6 PBM docks in the canonical binding groove of Scribble PDZ1 constituted by the β2 strand and helix α2. Superimposition of Scribble PDZ1 in complex with HPV16 E6 peptide with peptide free Scribble PDZ1(PDB ID 5VWC) [30] shows that HPV16 E6 does not influence the overall structure of Scribble PDZ, with a r.m.s.d of 1.2 Å over 96 Cα atoms [31].

Scribble PDZ1 binding to HPV16 E6 is achieved through the docking of L^0-HPV16^ in a PDZ1 pocket formed by PDZ1 L738, I740, I742, V796 and L800 (Figure 2A) as previously observed in other PDZ1 complexes [30, 32]. An ionic interaction between the guanidinium group of R^−^ 7-HPV16 and the side chain of E756^PDZ1^ support complex formation. A network of hydrogen bonds along the binding groove are also present, including the main chain mediated T^−2-HPV16^:I742^PDZ1^, L^0-HPV16^:I740^PDZ1^, L^0-HPV16^:L738^PDZ1^, L^0-HPV16^:G739^PDZ1^, R^−4-HPV16^:G744^PDZ1^ and Q^−1-HPV16^:I740^PDZ1^. Side chain mediated hydrogen bonds are observed between R^−4-HPV16^:S748^PDZ1^, E^−3-HPV16^:S761^PDZ1^, Q^−1-HPV16^:S741^PDZ1^ and T^−2-HPV16^:H793^PDZ1^. There were two PDZ1:HPV16 complexes in the asymmetric unit and most of the interactions described above are conserved between the two, with the E6 PBM peptide conformation highly conserved (Figure 2) with 0.4 r.m.s.d over the entire peptide chain.

Although the crystal structure of PDZ1:HPV18 closely resembles the PDZ1:HPV16 complex, several interactions have changed (Figure 2B). Q^−7-HPV18^ is involved in hydrogen bond with E756^PDZ1^, and the main chain of R^−6-HPV18^ forms a hydrogen bond with the main chain of T749^PDZ1^. The sequence of HPV18 differs from HPV 16 in the −6 position (T^−6^ in HPV16 and R^−6^ in HPV18), however HPV18 lacks this ionic interaction, though the interaction is maintained in the form of a hydrogen bond, between Q^−7^ with E756 ^PDZ1^. HPV 18 makes an additional ionic contact via R^−9-HPV18^ and E792 ^PDZ1^ in one of the two copies of the complex in the asymmetric unit.

Scribble PDZ3 binding to HPV16 E6 involves engagement of L^0-HPV18^ with the PDZ3 pocket formed by L1014^PDZ3^, L1016^PDZ3^ and L1078^PDZ3^ (Figure 2C). A hydrogen bond between R^−7-HPV16^ and the main chain of D1023^PDZ3^ is present. A network of hydrogen bonds is located along the binding groove, including the carboxyl group of L^0-HPV16^ contacts with L1014^PDZ3^ and G1015^PDZ3^ as well as main chain mediated T^−2HPV16^:I1018^PDZ3^, L^0-^ HPV16:L1016^PDZ3^ and T^−6-HPV16^:S1025^PDZ3^. Side chain mediated hydrogen bonds include T^−2-HPV16^:H1071^PDZ3^, E^-3-HPV16^:S1039^PDZ3^, R^-4-HPV16^:Q1072^PDZ3^ and T^-6-HPV16^:H1024^PDZ3^.

In the PDZ3:HPV66 complex (Figure 2D), V^0-HPV66^ is located in the PDZ3 pocket formed by L1014^PDZ3^, L1016^PDZ3^ and L1078^PDZ3^. The carboxyl group of V° forms main chain hydrogen bonds with L1014^PDZ3^, G1015^PDZ3^ and L1016^PDZ3^. Additionally hydrogen bonds are observed between S^−2-HPV66^ with the main chain amide of I1128^PDZ3^, and the imidazole ring of H1071^PDZ3^ via the Ser side chain hydroxyl group, as well as E^−3-HPV66^:S1039^PDZ3^, T^−1-HPV66^:S1017^PDZ3^ and T^-4-HPV66^:H1024^PDZ3^.

## DISCUSSION

It has previously been shown that E6 proteins from carcinogenic HPV strains are able to bind to key host proteins including the polarity regulators human Scribble and Dlg in order to subvert polarity signalling, and establish a cellular environment more supportive of viral infection [23]. However, the detailed molecular basis of the E6 interactions with Scribble has not been determined to date, although a significant role for the Scribble PDZ3 domain has been proposed as the main determinant for E6 interactions [25]. We systematically examined interactions of E6 from 7 carcinogenic and one non-carcinogenic HPV strain with individual Scribble PDZ domains to map in detail the interaction patterns underlying E6 subversion of host Scribble signalling. We then determined crystal structures of select interaction pairs including complexes of Scribble PDZ1 bound to E6 from HPV 16 and 18 as well as Scribble PDZ3 bound to E6 from HPB66. Our findings reveal distinct interaction patterns for each E6 PBM motif from a given HPV strain, suggesting that a complex molecular interplay exists that underpins the overt Scribble-HPV E6 interaction.

We found no direct correlation between measured affinity of an interacting HPV E6 PBM and its suggested carcinogenicity. HPV16 is reported to be the most carcinogenic of the strains, however the affinities of its E6 PBM motif compared to the less carcinogenic HPV strains are comparable and only differ by 2-3 fold. Interestingly, HPV16 E6 PBM did not interact with Scribble PDZ, as did HPV 66, whereas all other E6 PBMs originating from strains that showed an interaction with Scribble bound PDZ1, 2 and 3. HPV E6 is one of a number of virus encoded proteins that have been shown to bind to Scribble PDZ domains. Others include TBEV NS5, which binds Scribble PDZ3 with 18 μM affinity [14], avian influenza A virus Vietnam NS1 that binds Scribble to all four Scribble PDZ domains with affinities from 12-21 μM [15] and HTLV Tax [13], which also binds all four Scribble PDZ domains with affinities from 8-40 μM. For NS1 and Tax, the PDZ1 domain of Scribble is the highest affinity interactor, whilst PDZ4 interactions show the lowest affinities. Despite minor variations, the observed affinities of the identified PBMs for Scribble PDZ domains are comparable. Indeed, other PDZ-viral PBM interactions from other polarity regulatory proteins show similar affinities, including PALS1 PDZ:Sars-CoV2 E PBM interaction (23 μM affinity)[12].

When considering endogenous cellular interactions, HPV16 E6 peptide binds to Scribble PDZ1 with 9.4 μM affinity, which is similar to the affinity previously observed for endogenous interactor MCC [32]. Inspection and comparison of PDZ1:HPV16 E6 and PDZ1:MCC with high affinity complexes such as PDZ1:β-PIX [30] reveals the absence of hydrophobic side chains from the PBM engaging the PDZ1 β2-3 loop, as opposed to what is observed the in PDZ1:β-PIX structure. W641^*β*-PIX^ and W303^Erbin^ are present in position −4 of both high affinity complexes, which are occupied by R153 in HPV16 E6 (Figure 2C). In *D. melanogaster* PDZ1:GukH complex [33] the high affinity is achieved through a different mechanism where F1784^GukH^ forms a π-stacking interactions with H796^PDZ1^, but that interaction is not conserved in any other known Scribble interaction. Instead, in other Scribble PDZ domain complexes F1784^GukH^ is replaced by D642^β-PIX^, N825^MCC^ and R154^HPV16^ (Figure 2C). No comparable use of a hydrophobic side chain has been seen in interactions of viral PBM sequences with Scribble PDZ domains, in agreement with the reported weaker affinities [14, 15].

In addition to binding Scribble, HPV E6 PBMs have also been shown to bind to human Dlg1 and human phosphatases PTPN3 and 4. Affinity measurements of these interactions reveal that E6 PBM encoded by HPV51 binds to Dlg1 PDZ2 with affinities of 9.6 μM (11-mer E6 peptide) or 28.3 μM (6-mer E6 peptide) [34], whilst HPV16 and HPV18 E6 PBM bound PTPN3 with 53 μM and 37 mM affinity [35], respectively, whereas HPV16 E6 PBM bound PTPN4 with 19 μM affinity [36]. A structural analysis of the Dlg1 PDZ2:HPV51 E6 zinc binding domain complex (Figure 2F) [34] revealed complex formation is driven by a dense network of interactions. Notably, Dlg1 PDZ2 utilizes the side chain of K384 to contact the carboxyl group of the interacting PBM via an ionic interaction, which is not seen in our Scribble PDZ complexes where the equivalent residues are R801 and L1079/R1080 (PDZ1 and PDZ3). Furthermore, Dlg1 PDZ2 H384 is not involved in the canonical interaction with HPV 51 T-2. Whilst the overall mode of binding of Scribble and Dlg PDZ domains with HPV E6 PBM is conserved, clear differences in the detailed interaction exist, although they do not manifest themselves as substantial drivers for large affinity differences between the interactions.

In summary, we examined the biochemical and structural basis for HPV E6 interactions with the human cell polarity regulator Scribble. Our findings suggest that distinct interaction patterns for each E6 PBM motif from a given HPV strain exist, and reveal a complex molecular interaction network between Scribble and HPV E6 that influences E6 carcinogenic potential. Furthermore, our data may serve as a platform for the design of novel antiviral therapeutics targeting the HPV E6 – Scribble interaction [37].

## METHODS

### Protein expression and purification

Protein expression constructs encoding the PDZ domains of human Scrib (Uniprot accession number: Q14160) PDZ1 (728–815); PDZ2 (833–965); PDZ3 (1005–1094); and PDZ4 (1099–1203)) were obtained as synthetic cDNA codon optimized for *Escherichia coli* expression and cloned into the pGil-MBP [38] and pGex-6P3 (GE Healthcare) as previously described [30]. Protein overexpression was performed using *Escherichia coli* BL21 (DE3) pLysS cells (BIOLINE) in super broth supplemented with 200 μg/mL ampicillin (AMRESCO) using auto-induction media (10 mM Tris-Cl pH7.6, 100 mM NaCl, 1 mM MgSO4, 0.2 % (w/v) D-lactose, 0.05 % (w/v) glucose, 0.5 % (v/v) glycerol) [39] at 37°C until the optical density at 600 nm (OD600) reached 1.0 before transferring cultures to 20°C for 24 hours for protein expression. Target Scribble PDZ domains were purified as previously described [30]. All target proteins were subjected to size exclusion chromatography using the HiLoad 16/600 Superdex 75 (GE Healthcare) equilibrated in 25 mM Tris pH8.0, 150 mM NaCl, and eluted as a single peak.

### Isothermal titration calorimetry

Purified human Scribble PDZ domains were used in titration experiments against 10-mer peptides spanning the C-terminus of different strains of human papilloma virus (HPV) E6 protein to measure their affinity for Scribble PDZ domains. E6 peptides used were from HPV16 (Uniprot accession number: Q919B4, SSRTRRETQL), HPV18 (Uniprot accession number: P06463, RLQRRRETQV), HPV31 (Uniprot accession number: P17386, WRRPRTETQV), HPV33 (Uniprot accession number: P06427, WRSRRRETAL), HPV40 (Uniprot accession number: P36812, EKGQRSETLC), HPV51 (Uniprot accession number: P26554, RTRQRNETQV), HPV66 (Uniprot accession number: Q80955, TSRQATESTV) and HPV70 (Uniprot accession number: P50804, RRRIRRETQV).

PDZ1 concentration was quantitated at 280 nm absorbance (A280nm) using a NanoDrop 2000/2000c UV-Vis Spectrophotometer (Thermo Scientific). Since the concentration of PDZ2, PDZ3 and PDZ4 could not be determined by measurements of Abs280nm due to lack of useful aromatic amino acids, protein concentrations were calculated using the Scope method [40] by measuring absorbance at 205 and 280 nm using a NanoDrop 2000 spectrophotometer (Thermo Fisher Scientific).

Titrations were performed at 25°C with a stirring speed of 750 rpm using the MicroCalTM iTC200 System (GE Healthcare) as previously described [41]. A protein concentration of 75 μM against peptide concentration of 0.9 mM were used. Peptides were purchased from Mimotopes (Mulgrave, Australia). Raw thermograms were processed with MicroCal Origin® version 7.0 software (OriginLabTM Corporation) to obtain the binding parameters of each interaction. A synthetic pan-PDZ binding peptide referred to as superpeptide (RSWFETWV) was used as a positive control [42].

### Protein crystallisation, data collection and refinement

Complexes of PDZ1 and PDZ3 with HPV E6 PBM peptides were reconstituted by mixing protein and peptide at a 1:2 molar ratio as previously described [41]. Crystallization trials were carried out using 96-well sitting-drop trays (Swissci) and vapor diffusion at 20 °C either in-house or at the CSIRO C3 Collaborative Crystallization Centre, Melbourne, Australia. 0.15-μl of protein-peptide complexes were mixed with 0.15 μl of various crystallization conditions using a Phoenix nanodispenser robot (Art Robbins). Commercially available screening kits (PACT Suite and JCSG-plus Screen) were used for the initial crystallization screening, with hit optimization performed using a 96-well plate at the CSIRO C3 Centre. Crystals of PDZ1 in complex with HPV16 E6 peptide was obtained at 23 mg/ml in 20% (w/v) polyethylene glycol 6000, 0.2 M ammonium chloride and 0.1 M HEPES pH 7.0. The PDZ1-HPV16 crystals were cryo-protected using 20% (w/v) glucose and flash-cooled at 100 K using liquid nitrogen. Plate shaped crystals were obtained belonging to space group C2. Crystals of PDZ1 in complex with HPV18 E6 peptide were obtained at 12 mg/ml in 12% w/v polyethylene glycol 4000 and 0.1 M ammonium sulfate. The PDZ1-HPV18 crystals were cryoprotected using 25% ethylene glycol and flash-cooled at 100 K using liquid nitrogen. Rod shaped crystals were obtained belonging to the space group P2_1_2_1_2_1_.

Crystals of PDZ3 in complex with HPV66 E6 peptide were obtained at 10 mg/ml1 in 63% (v/v) MPD and 25 mM HEPEs pH 7.5. The PDZ3-HPV66 crystals were cryoprotected using 20% (w/v) glucose and flash cooled at 100 K using liquid nitrogen. Diamond plate shaped crystals were obtained belonging to the space group C2. Crystals of PDZ3 in complex with HPV16 E6 peptide were obtained at 10 mg/ml in 2.5 M NaCl and 0.1 M HEPES pH 8.0. The PDZ3-HPV16 crystals were cryoprotected in 25% 2-propanol and flash cooled at 100 K using liquid nitrogen. Rhombohedral crystals were obtained belonging to the space group P6_1_22.

Diffraction data were collected on the MX2 beamline at the Australian Synchrotron using an Eiger 16M detector with an oscillation range of 0.1° per frame using a wavelength of 0.9537 Å. Diffraction data were integrated with Xia2 [43] and scaled using AIMLESS [44]. Structures was solved by molecular replacement using Phaser [45] with the structure of human Scribble PDZ1 (PDB code 5VWC [30] or human Scribble PDZ3 (PDB code 5VWI [30]) as search models. The solutions produced by Phaser was manually rebuilt over multiple cycles using Coot [46] and refined using PHENIX [47]. Data collection and refinement statistics details are summarized in Table 2. MolProbity scores were obtained from the MolProbity web server [48]. Coordinate files have been deposited in the Protein Data Bank under the accession codes 8B87 (PDZ1:HPV16), 8B82 (PDZ1:HPV18), 8B8O (PDZ3:HPV66) and 8B9T (PDZX3:HPV66). All images were generated using the PyMOL Molecular Graphics System, Version 2.5 Schrödinger, LLC. All software was accessed using the SBGrid suite [49]. All raw diffraction images were deposited on the SBGrid Data Bank [50] using their PDB accession 8B87 (PDZ1:HPV16), 8B82 (PDZ1:HPV18), 8B8O (PDZ3:HPV66) and 8B9T (PDZX3:HPV66).

## Author Contributions

BZS: Acquisition of data; Analysis and interpretation of data. Drafting and revising the article.

SC: Acquisition of data; Analysis and interpretation of data; Drafting and revising the article.

POH: Conception and design; Analysis and interpretation of data; Drafting and revising the article.

MK: Conception and design; Acquisition of data; Analysis and interpretation of data; Drafting and revising the article.

## Acknowledgements

We thank staff at the MX beamlines at the Australian Synchrotron for help with X-ray data collection, and the CSIRO C3 Collaborative Crystallization Centre for assistance with crystallization and the Comprehensive Proteomics Platform at La Trobe University for core instrument support. We thank the ACRF for their support of the Eiger MX detector at the Australian Synchrotron MX2 beamline. This work was supported in whole or part by the National Health and Medical Research Council Australia (Project Grant APP1103871 to MK, POH; Senior Research Fellowship APP1079133 to POH), Australian Research Council (Fellowship FT130101349 to MK) and La Trobe University (Research focus area “Understanding Disease” project grant and scholarship to BZS). The funders had no involvement in this study.

